# The changes in the p53 protein across the animal kingdom pointing to its involvement in longevity

**DOI:** 10.1101/2020.05.06.080200

**Authors:** Martin Bartas, Václav Brázda, Adriana Volná, Jiří Červeň, Petr Pečinka, Joanna Zawacka-Pankau

**Affiliations:** Department of Biology and Ecology, Faculty of Science, University of Ostrava, Ostrava, 710 00, Czech Republic; Institute of Biophysics, Czech Academy of Sciences, Brno, 612 65, Czech Republic; Department of Physics, Faculty of Science, University of Ostrava, Chittussiho 10, Ostrava, CZ, 71000, Czech Republic; Faculty of Chemistry, Pasteura 1, 02-093 Warsaw, University of Warsaw, Warsaw, Poland; &Department of Medicine, Huddinge, Center for Hematology and Regenerative Medicine, Karolinska Institute, Stockholm, SE 171-74 Sweden

**Keywords:** p53, aging, longevity, bioinformatics, sequence analysis

## Abstract

Recently, the quest for the mythical fountain of youth has turned into extensive research programs aiming to extend the healthy lifespan in humans. Despite advances in our understanding of the aging process, the surprisingly extended lifespan and cancer resistance of some animal species remains unexplained. The p53 protein plays a crucial role in tumor suppression and in tissue homeostasis and aging. Long-lived, cancer-free African elephants, have 20 copies of TP53 gene including 19 retrogenes (38 alleles) which are partially active, whereas humans possess only one copy of TP53 and have an estimated cancer mortality of 11-25%. The mechanism through which p53 contributes to the resolution of the Peto’s paradox in the Animalia remains vague. Thus, in this work, we took advantage of the available datasets and inspected the p53 amino acid sequence of phylogenetically related organisms that show variations in the lifespan. We discovered new correlations between specific amino acid deviations in p53 and the lifespans across different animal species. We found that species with extended lifespan have certain characteristic amino acid substitutions in the p53 DNA binding domain that alter its function as depicted from the Phenotypic Annotation of p53 Mutations, using PROVEAN tool or SWISS-MODEL workflow. Our findings show a direct association between specific amino acid residues in p53 protein, changes in p53 functionality and the extended animal lifespan, and further highlight the importance of p53 protein in aging.

## 1. Introduction

The promise of the eternal life have inspired research into this topic across many civilizations and through the millennia dating back to Herodotus and his writings 2,500 years ago. Although the average human lifespan is increasing, our health span appears to be lagging. Several studies argue that human lifespan is physiologically and genetically limited [1], yet recent contributions have proposed a future, potentially unlimited increase of human lifespan [2]. The demographical data show that the death risk increases exponentially up to about age 80, then decelerates and plateaus after age 105 [3]. There are two major theories of aging, senescence theory and programmed theory of aging [4]. The senescence theory converges on the accumulation of the cellular damage that cannot be repaired leading first to the permanent cell-cycle arrest and in the end loss of the organismal fitness. The free radical theory, can be classified as a subtype of senescence theory and postulates that organisms age because of the accumulation of damage inflicted by reactive oxygen species [5,6]. There is also a common agreement that the preservation in the fidelity of the DNA repair process involving the p53 pathway favors longevity [7]. The programmed theory of aging states that ageing is tightly controlled and includes Hayflick limit theory and the central aging clock theory. At the molecular level, the biological aging is a complex process that involves genetic factors, mitochondria-damage mechanisms, cellular senescence, proteostasis and autophagy, telomeres attrition, epigenetics, inflamation and metabolic switch. Thus, the lifespan is a multi-nodal characteristic [8]. To date, several proteins have been found to play important roles in human aging including mTOR, AMPK, SIRT1, PGC1α, APOE, LPA, CDKN2B-AS1 and p53. Among those, p53 emerges as a central node, linking several pathways together. The p53 is a tumor suppressor coded by the most often mutated gene in human cancers [9–12], and the loss of wild-type p53 function is associated with the fatal outcomes in cancer patients. p53 is a critical sensor of cellular stress and the dictator of the cell fate. Depending of the types of stress which include DNA damage, oncogene activation, nutrient deprivation, reactive oxygen species accumulation or telomere shortening, p53 either transiently stops cell proliferation and initiates the DNA repair machinery, induces cell death or pushes cells to replicative senescence, a permanent proliferation arrest.

Taken high cancer susceptibility in humans and the role of p53 in regulating cell fate in health and disease, p53 is regarded as the key regulator of human healthy lifespan [13,14]. When we consider the “lifespan” of tumor cells, it is apparent that cancer cells often gain new functions, including “immortality”, at least partially attributed to the mutations in the *TP*53 gene and/or in its regulatory pathways [15]. As reviewed by Stiewe and Haran [16], cancer-associated mutations alter p53 in three ways: they promote loss of wild-type (wt) p53-DNA binding, trigger dominant-negative inhibition of wtp53 by the mutant p53 in the monoallelic mutation setting, or induce gain of new functions to mutant p53 through new protein-protein-DNA interactions. Loss of binding to the canonical target sequnece by mutant p53 can be partial or complete. Different mutant p53 proteins show a variable degree of loss of the DNA binding capacity. This results in the attenuated or target-selective DNA binding patterns [16]. Multiple functions of p53 have been described and extensively reviewed [17–19]. For example, the p53 protein plays roles in metabolism [20], cell cycle arrest [21,22], apoptosis [23], ferroptosis, angiogenesis [24], DNA repair [25], embryonic development and cell senescence [18,26]. In the majority of the cellular processess, p53 functions as a transcription factor and recognizes and binds to multiple target genes through a recognition sequence (5’-PuPuPuC(A/T)(T/A)GPyPyPy-3’) [27–29]. Owing to its crucial role in protection against the accumulation of DNA damage, p53 is called “the guardian of the genome” [30,31].

From the evolutionary point of view, the *TP*53 gene is specific for the Holozoa branch and its ancestral p63/p73-like genes emerged approximately one billion years ago [32,33]. The p53/p63/p73 family plays key roles in several major molecular and biological processes including tumor suppression, fertility, mammalian embryonic development and aging [20]. Unlike *TP*53, *TP*73 and *TP*63 genes are rarely mutated in cancers. Yet, the tumor suppressor function of p73 is often attenuated in humans. The mechanism of suppression is by hypermethylation of CpG islands at promoter 1, the binding to the overexpressed dominant negative isoform, dNp73 [34] or to MDM2 or MDMX. The pharmacological inhibition of protein-protein interactions is currently explored for the improved cancer therapy [35]. Notably, it has been demonstrated that all p53 family members take part in regulating aging through activation of senescence and regulating DNA repair [18,25].

p53 is a major factor regulating cellular senescence and the mechanism is by activation of CDKN1A (p21) and PML. The study by Tyner *et al*., showed that heterozygous mice having one *TP*53 allele with deletion of the first six exons (p53^+/m^, Δ exon 1-6) are aging prematurely. This mutant mice exhibited enhanced resistance to spontaneous tumors, yet displayed accelerated aging compared to p53^+/+^ mice [36]. The study by same group showed that truncated p53 protein stabilized wild-type p53 in non-stressed cells, promoted its nuclear accumulation and induced hyper-stability of wild-type p53 upon irradiation [37]. Based on this observation the conclusion was made that the constitutive expression of p53 accelerates aging. This was not confirmed in a follow-up study [38], as the pro-aging phenotype was not seen in the p53 “super-mice”, expressing additional copies of *TP*53 gene. Thus, over-activated p53 *per se* might not be a critical driver of accelerated aging. Yet, the role of p53’s hyper-activity in aging appears to be conflicting. Fibroblasts derived from hereditary segmental progeroid syndrome patients with the homozygous antiterminating mutation, c.1492T>C, in *MDM*2 gene, showed p53 hyperstability and accelerated aging [39]. This study postulates that hyper-stability of p53 due to abberant MDM2-p53 axis and the exposure to chronic stress induces the aging phenotype through the induction of chronic senescence. MDM2, the best-described negative regulator of p53, binds to the N terminal domain of p53 by its N-terminus. The knockout *Mdm*2−/− mice are embryonic lethal in wtp53 background which indicates that p53 regulation by MDM2 is critical for development. Yet, the conditional deletion of *Mdm*2 in the epidermis induced p53-mediated senescence and accelerated aging [40]. Thus, deregulated MDM2-p53 axis might play a role in aging phenotype.

Gradual DNA damage and mitochondrial decline are hallmarks of physiological aging. DNA damage activated by the telomere attrition in an aging cell induces p53 and mitochondrial dysfunction through repression of the PPARγ co-activator 1α (PGC1α). This induces senescence [41]. Also, the study on the hereditary segmental progeroid syndrome clearly manifested the role of MDM2 inactivation and p53 hyperactivity in the aging phenotype [39]. Despite the emerging evidence, the exact molecular mechanisms underlying the p53-mediated aging phenotype need to be elucidated. For example, it has been demonstrated that replicative senescence is facilitated by p53 mainly through activation of CDKN1A/p21. Yet, there are several other factors contributing to aging like activation of E2F and mTOR as described elsewhere [18]. In principle, it can be concluded that p53 prevents cancer and protects from aging under physiological conditions, however, chronic stress-amplified p53 has a detrimental effect on healthy aging despite retaining its tumor suppression function. Hence, p53 can either be a pro-aging or a pro-longevity factor, depending on the physiological context [42].

In addition to full-length p53, p53 isoforms may also play an important role in the modulation of longevity. The expression of certain short and long p53 isoforms might contribute to a balance between tumor suppression and tissue regeneration [43]. In example, p53β isoform, generated through the alternative splicing of intron 9, is upregulated in normal human senescent fibroblasts and interacts with full-lengts p53 to induce CDKN1A [44].

Considering the critical role of p53 in maintaining tissue homeostasis, high frequency of gain of function mutations in cancer and the limited and the conflicting information on p53 role in organismal aging in Animalia, in the present work we employed currently available datasets and tools and analyzed p53 protein sequences in species possessing an extended lifespan. Our thorough analysis depicted a suprising correlation between the changes in the p53 protein sequence and the organismal lifespan, both in short- and long-lived species. Many of the identified changes occurred in the DNA binding domain and might have a detrimental effect on p53 DNA-binding activity. All in all, we found that, when compared to the majority of closely related organisms within their phylogenetic groups, animals with unusually long lifespans share atypical p53 protein sequence features in the position corresponding to human 180-192 p53 region, pointing to the important contribution of changes in p53 in life expectancy.

## 2. Results

Since the growing evidence implies that p53 activity plays a pivotal role in aging in humans and little is known about the molecular signatures of extending lifespan in animals, we inspected all currently available sequence data of long-lived animals to explore a link between longevity (maximal lifespan) and p53 protein sequences. For this, we used the longevity data from the AnAge Database [45]. We merged all available p53 sequences from the RefSeq database with AnAge Database (for more detail refer to Materials and methods). The p53 sequence from 118 species and their lifespan data were catalogued and sorted according to their phylogenetic group (Supplementary Material 1).

The longest living animal in our dataset is the bowhead whale (*Balaena mysticetus*) from **Artiodactyla** (subgroup Cetacea) with a maximal lifespan of 211 ± 35 years [46]. Bowhead whales have a significantly longer lifespan (about four times longer) compared with other whales. A thorough comparison of p53 protein sequences showed that, in contrast to other Cetacea, *Balaena mysticetus* has a unique leucine substitution in the proline-rich region, corresponding to amino acid residue 77 in human p53 (Figure 1). Even though, the change in the amino acids is predicted to be neutral according to the Protein Variation Effect Analyzer (PROVEAN) score of -0.993, the substitution still might change the activity of p53. Yet, this could only be addressed by the extended functional studies. All other accessible p53 sequences of whales have identical amino acid residue in this position as human p53.

**Figure 1.**
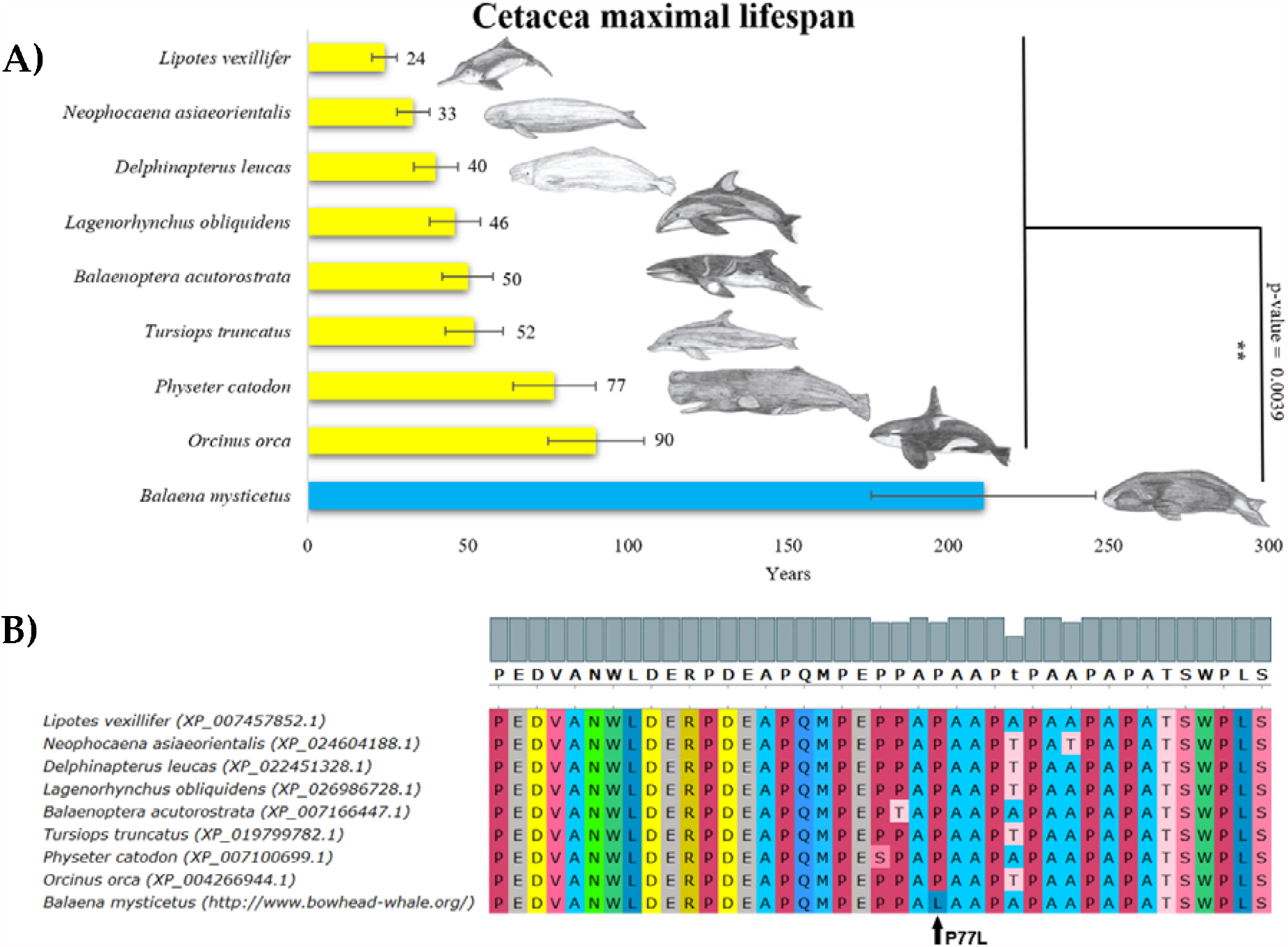
Cetacea lifespan and the corresponding p53 sequence changes. (A) Comparison of maximal cetacea lifespan in years. Bowhead whale (*Baleana mysticetus*) maximal lifespan is more than twice the maximal lifespan of the rest of cetacea (Wilcoxon one-sided signed-rank test was used, p-value < 0.05). (B) Multiple sequence protein alignments of p53 proline-rich region, performed in MUSCLE with default parameters [47], colors in “UGENE” style.

Most amphibian species live for less than 30 years [45], however, olm (*Proteus anguinus*, Batrachia, Amphibians), the only exclusively cave-dwelling chordate, has a maximal documented lifespan of 102 years. Comparison of the p53 protein sequences in amphibians showed a previously unrecognized insertion in *Proteus anguinus*. The p53 protein from this species has additional serine and arginine residues in its core domain (corresponding to insertion after amino acid L188 in human p53) which according to PROVEAN tool has the deleterious effect on p53 functionality (Figure 2, Table 1).

**Figure 2:**
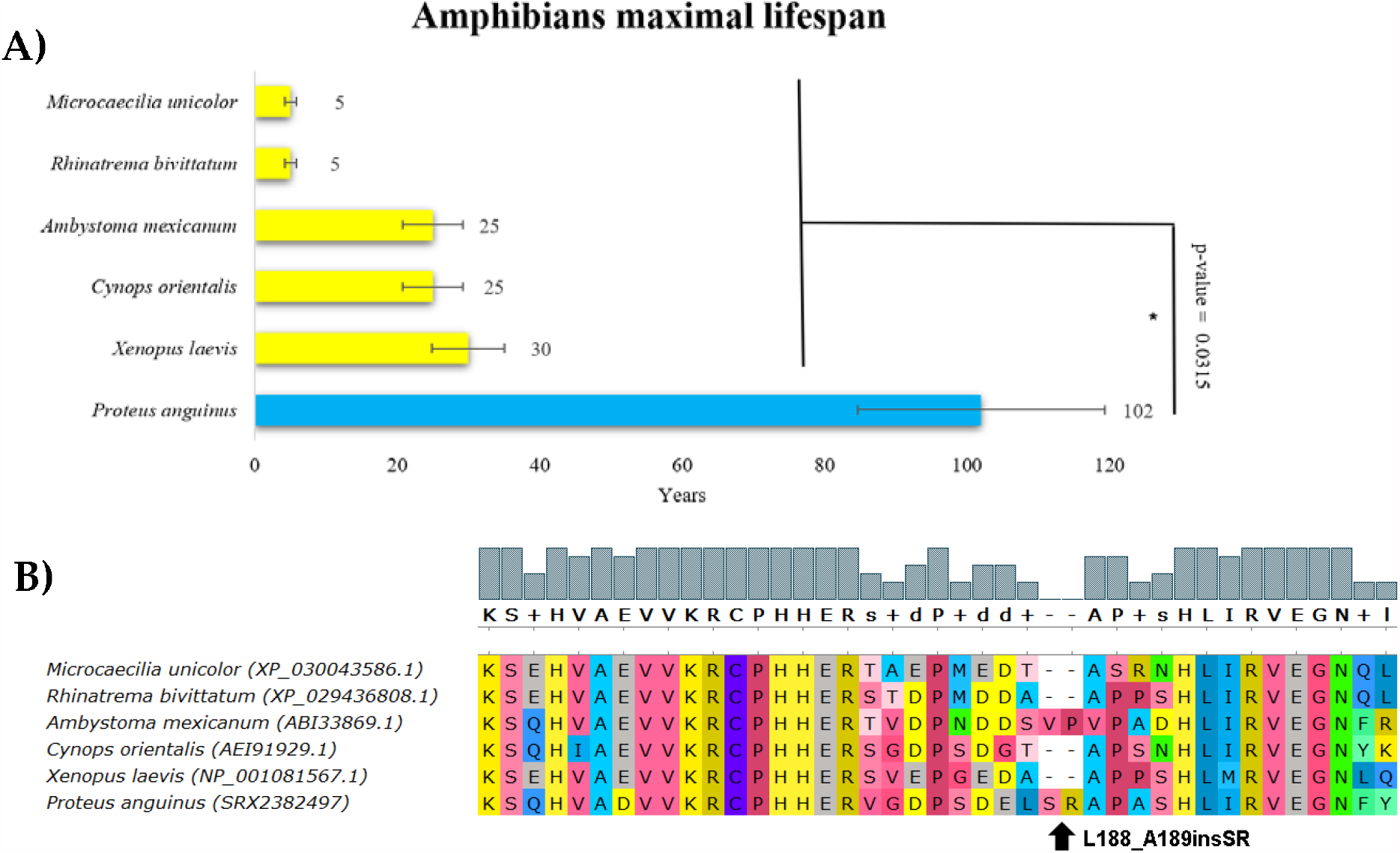
Amphibians lifespan and the corresponding p53 sequence changes. (A) Comparison of amphibian lifespans in years. Olm’s (*Proteus anguinus*) maximal lifespan is more than three times higher than the maximal lifespan of other amphibians (Wilcoxon one-sided signed rank test, p-value < 0.05). (B) Multiple protein alignments of the p53 dimerization region. Olm (*Proteus anguinus*) has a two-amino acid-residue long insertion following amino acid residue 188 (related to human p53 canonical sequence). The sequence of the p53 homolog from *Proteus anguinus* was determined using transcriptomic data from the SRA Archive (SRX2382497). Methods and color schemes are the same as in Figure 1B.

**Table 1:**
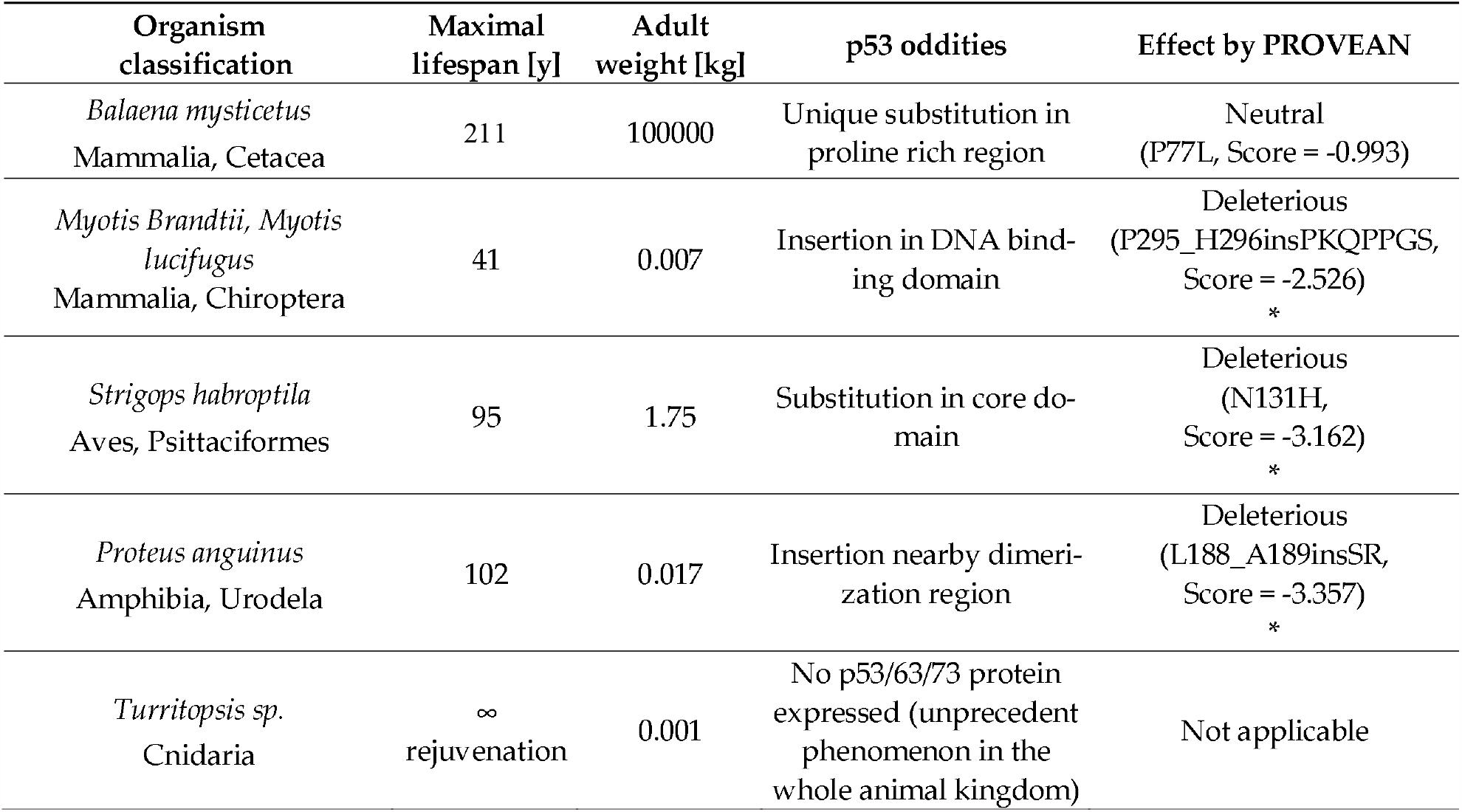
Comparison of animals characterized by extreme longevity and their atypical p53 features, where significance of particular changes was predicted. Default PROVEAN threshold -2.5 was used, insertions and deletions were submitted in respect to the human canonical protein sequence (NP_001119584.1) “*” indicates significant PROVEAN values (<-2.5).

Kakapo (*Strigops habroptila*, **Aves**) is a long-lived, large, flightless, nocturnal, ground-dwelling parrot endemic to New Zealand with a lifespan of around 95 years (Figure 3A, blue bar). Comparison of p53 protein sequence with other related species shows a change at positions 128 and 131, corresponding to the following change in human p53; P128V and N131H (Figure 3B). Interestingly, N131H mutations in human p53 are found in pancreatic and colon cancers [48,49]. This mutation most probably changes the structure of the p53 core domain and decreases the ability of p53 to bind to canonical DNA sequence. Relevantly, according to PHenotypic ANnotation of *TP*53 Mutations (PHANTM) classifier, the N131H mutation decreases p53 transcriptional activity by 47.19% [50]. In addition, according to the PROVEAN tool, substitutions at position 128 are deleterious with the score -4.45 (Table 1). These findings support the hypothesis that the change in p53 in kakapo is linked to the loss of function. We speculate that the lack of exposure to sunlight, thus low incidence of UV-induced DNA damage, might render p53 inactive in this species.

**Figure 3:**
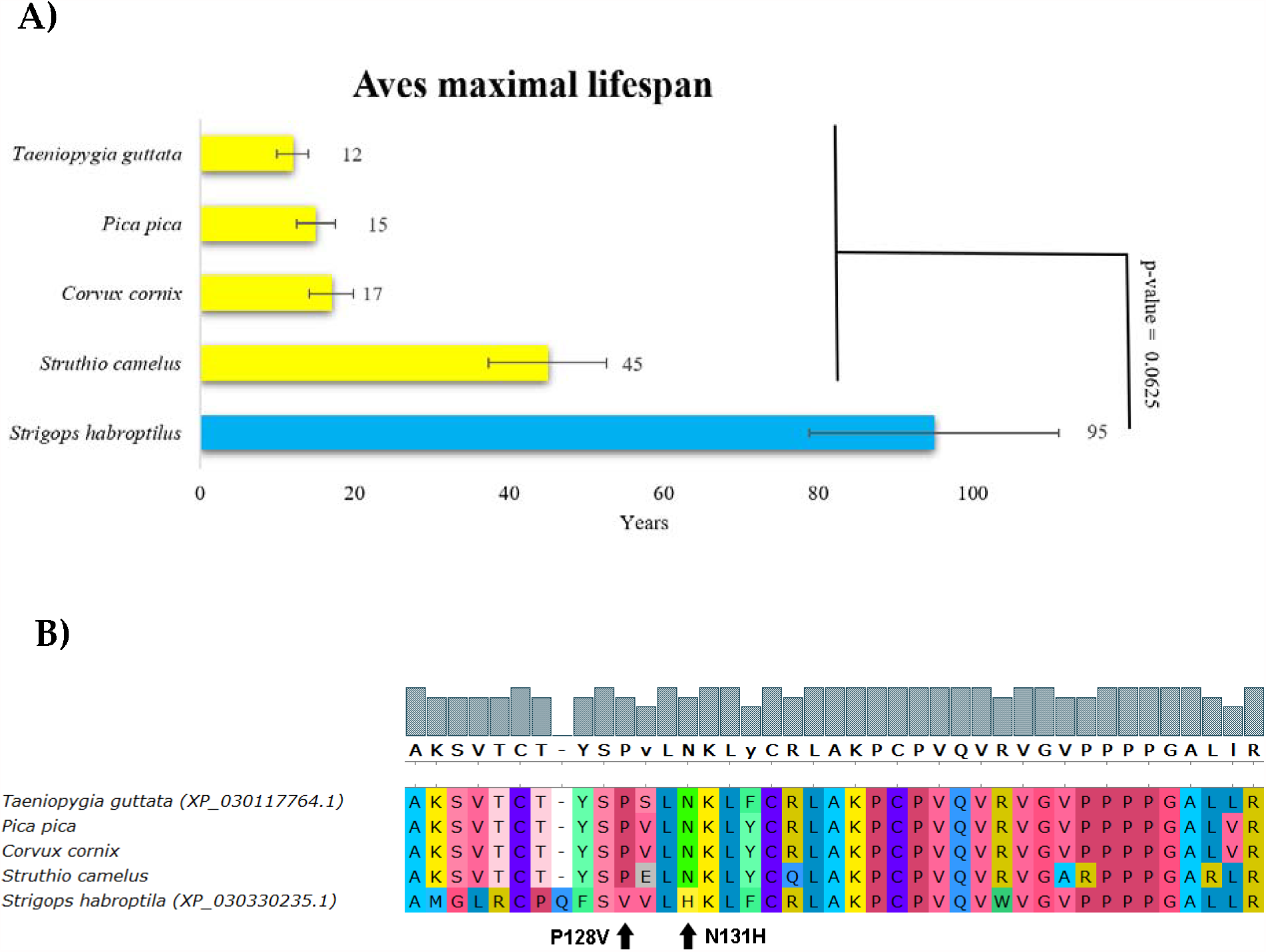
Aves lifespan and the corresponding p53 sequence changes. (A) Comparison of maximal Aves lifespan in years. Kakapo’s (*Strigops habroptila*) maximal lifespan is more than twice the maximal lifespan of other Aves (Wilcoxon one-sided signed rank test, p-value < 0.05). (B) Multiple protein alignments representing partial p53 core domain of the accessible Aves sequences. Sequences of all avian p53 homologs were determined using transcriptomic data from the SRA Archive, except from *Strigops habroptilus*, where the p53 sequence was known (XP_030330235.1). Methods and color schemes are the same as in Figure 1B.

Next, our analysis identified alterations in p53 in species having long lifespan in the **Chiroptera** group. The Brandt’s bat (*Myotis Brandtii*) is an extremely long-lived bat with a documented lifespan of 41 years [51]. Together with its close relative *Myotis lucifugus* they have significantly longer lifespans than other bats (Figure 4 A, blue bars). These two species share an unique arrangement in the p53 DNA binding region, with the seven amino acid residue insertion in the central DNA-binding region (following amino acid 295 in the human p53 canonical sequence) (Figure 4B). To assess how this rearrangement in the DNA binding region changes the interaction of p53 with DNA, we next modelled the p53 tetramer using SWISS-MODEL workflow. The insertion in the DNA binding domain of bats with long lifespan occurs in the DNA interaction cavity, suggesting decreased affinity of p53 for binding to DNA (Supplementary Material 2). Myotis Brandtii and Myotis lucifugus are very small bats (max 8 g bodyweight) and form a significant exception from the Max Kleiber’s law (mouse-to-elephant curve), since their lifespan is extremely long in relation to their small body size [52].

**Figure 4:**
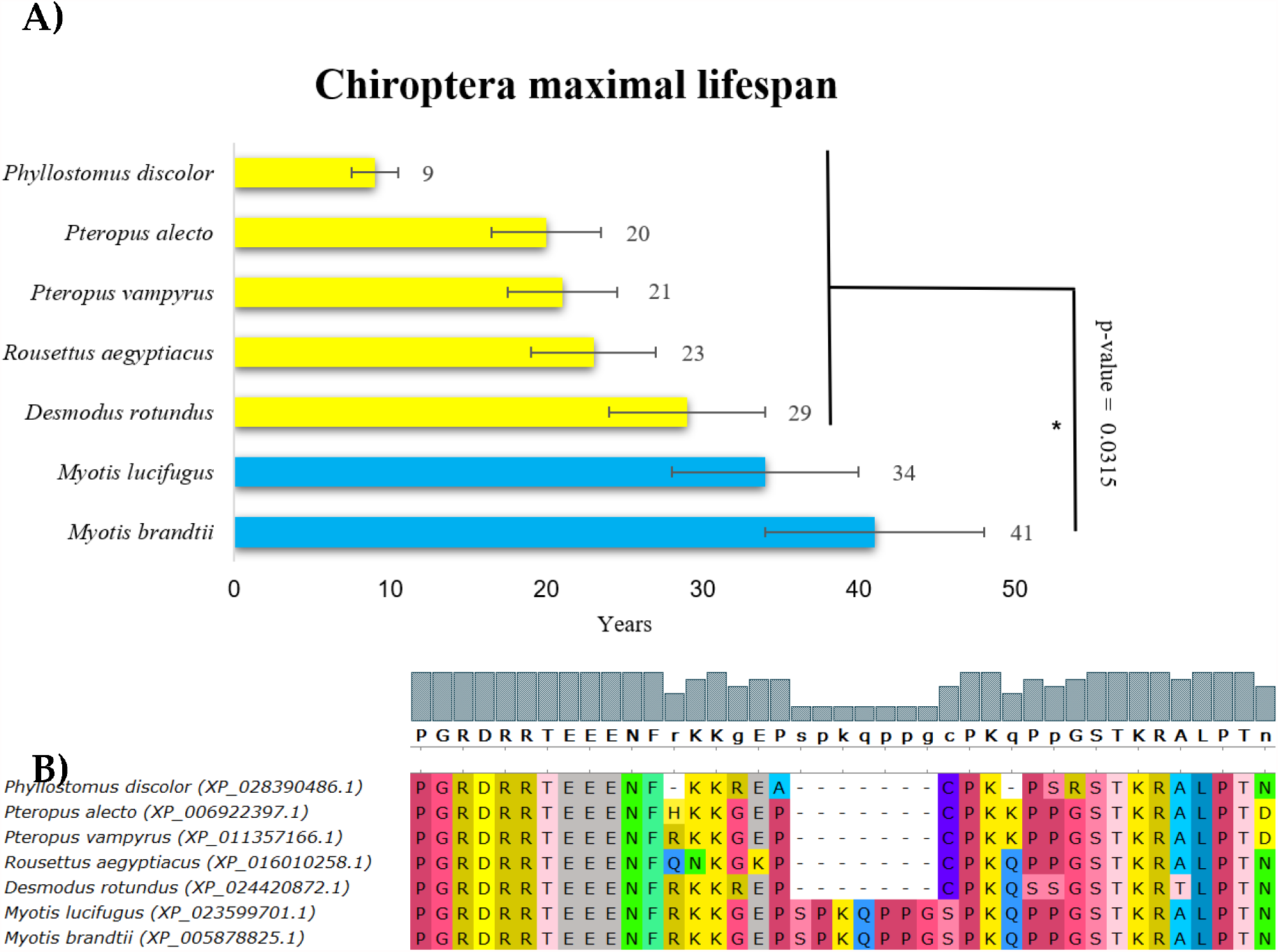
Chiroptera lifespan and the corresponding p53 sequence changes. (A) Comparison of maximal Chiroptera lifespan in years. Brandt’s bat (*Myotis brandtii*) and *Myotis lucifugus* maximal lifespans are significantly longer compared with other sequenced bats (Wilcoxon one-sided signed rank test, p-value < 0.05). (B). Multiple protein alignments of the C-terminal part of the p53 core domain of accessible Chiroptera sequences. Methods and color schemes are the same as in Figure 1B.

The abovementioned analysis of long-lived organisms in various animal groups led us to conclude that the amino acid sequence of p53 is associated with organismal lifespan. Therefore, we continued the analysis by further correlating the p53 amino acid sequence with the lifespan across the animal kingdom. Due to low similarity between p53 N-terminal and C-terminal domains across species and a significant role of mutations in the p53 DNA binding domain in cancer, we focused on the most conserved core domain of p53 and constructed the p53-based tree (Figure 5, left panel) [32]. We then compared contemporary phylogenetic tree with the tree based on p53 protein sequence (Figure 5). Then, the dataset with p53 sequences and animal lifespans were divided into 12 groups based on their phylogenetic relationships. Interestingly, some p53 sequences are not closely associated with the phylogenetic tree, indicating several parallel evolutionary processes leading to modified p53 activity. Even closely related species in various groups have significantly different lifespans (Supplementary Material 1) and therefore, are suitable for correlation analyses according to the method introduced by Jensen and colleagues [53].

**Figure 5:**
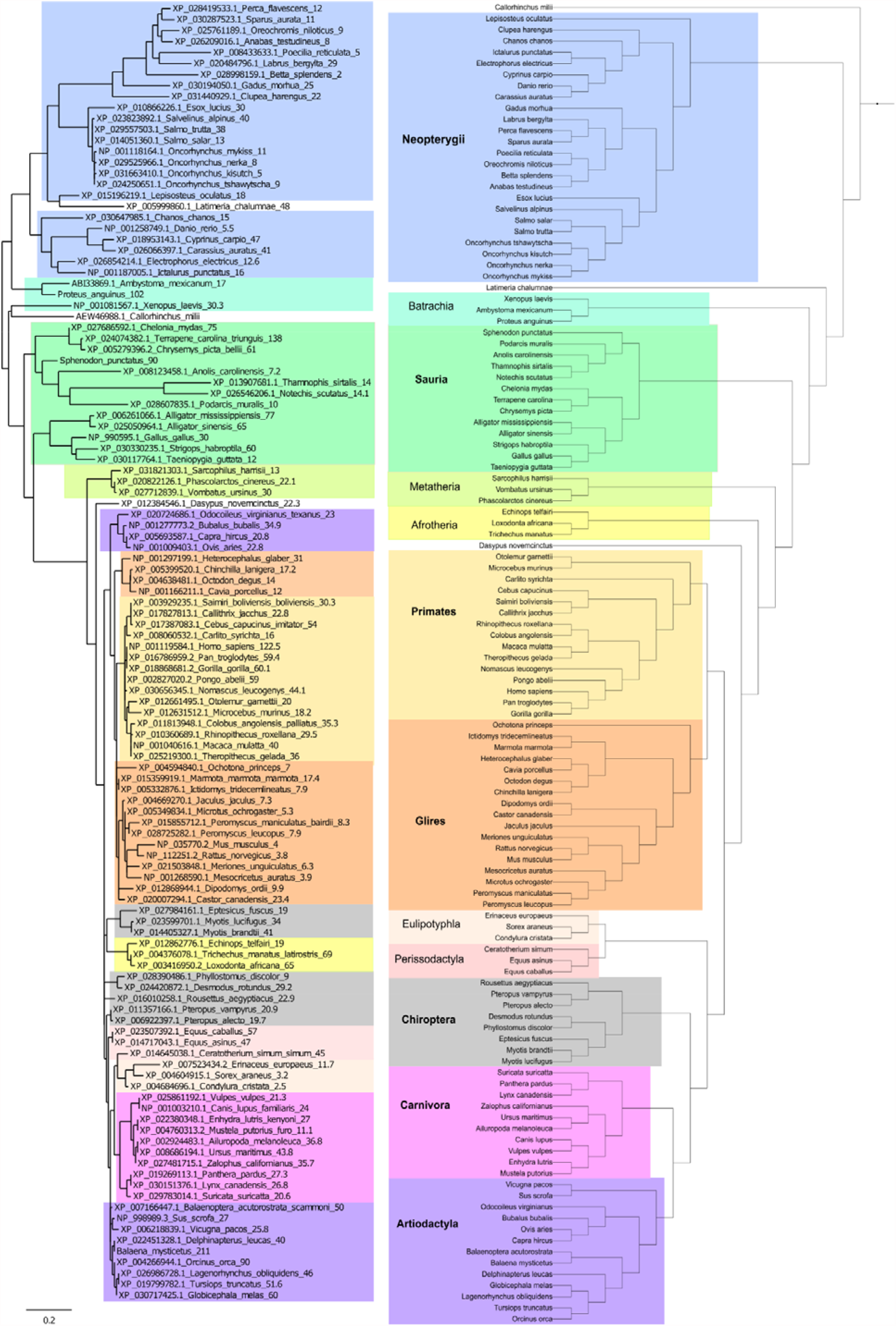
The p53-based and contemporary phylogenetic trees. Comparison of p53 protein tree (left) and the real phylogenetic tree (right). The protein tree was built using Phylogeny.fr platform. Organismal phylogeny was reconstructed using PhyloT and visualized in iTOL (see Methods for details). The color background represents the same phylogenetic groups.

Figure 6 summarizes the lifespan data and the total number of analyzed animals for each group with minimal and maximal values shown in Supplementary Material 3. Only datasets with more than 5 members in the group were used in the correlation analyses.

**Figure 6:**
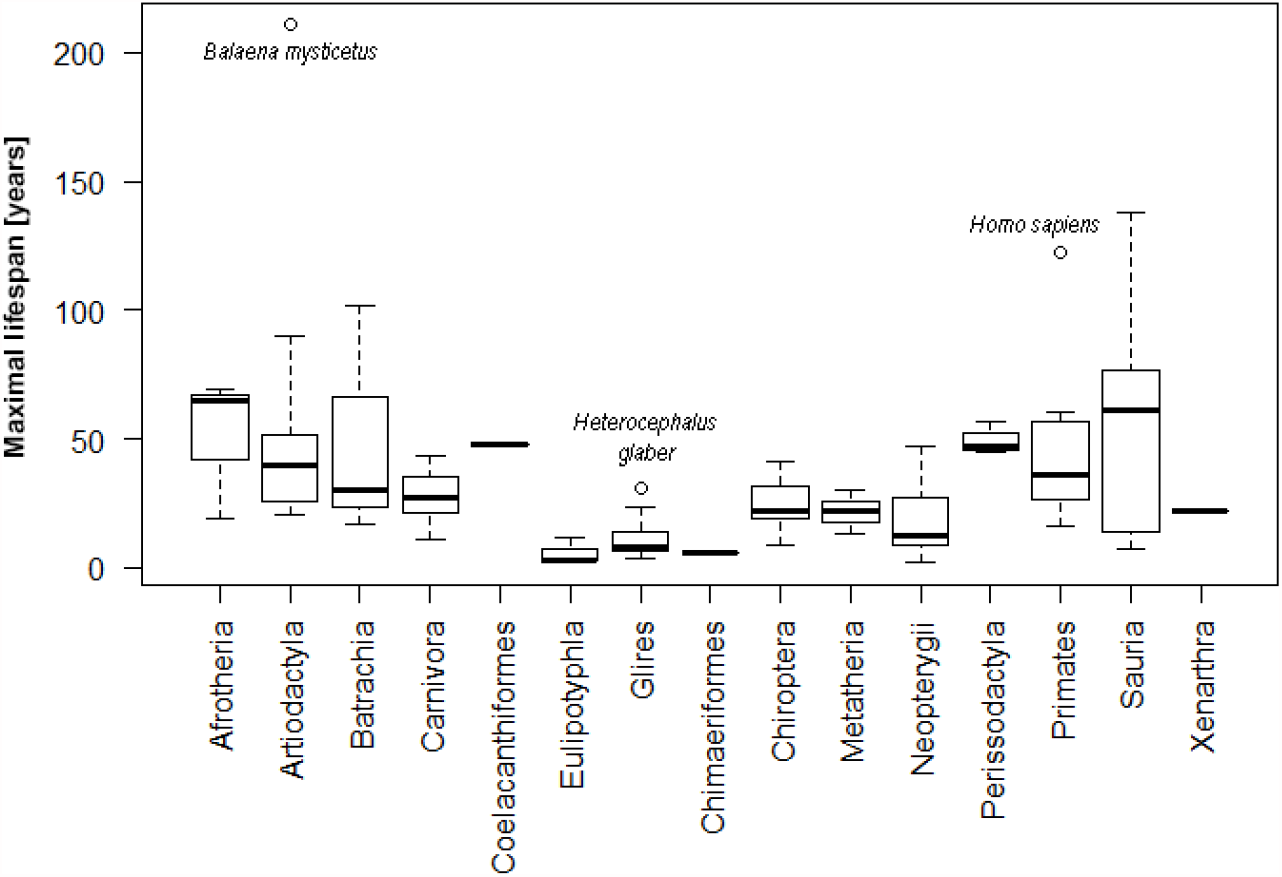
Representation of lifespans for all tested phylogenetic groups.

The organisms with the longest lifespan in the **Neopterygii** dataset are the carp (*Cyprinus carpio* (47 years)), followed by the goldfish (*Carassius auratus* (41 years)). Siamese fighting fish (*Betta splendens)* has the shortest lifespan in the group (2 years). The correlation analyses show that fifteen-amino acid-residues in the p53 core domain are significantly associated with prolonged lifespan (Figure 7A). We found that the most common variation in the long-lived Neopterygii is the presence of serine at positions corresponding to 98, 128 and 211 of human p53, and the presence of valine at positions 128, 150, 217, 232. On other hand, in the short-lived organisms in Neoropterygii, we identified threonine at positions 98, 100, 141, 217, 260, glutamic acid at positions 110, 128, 150 and 291 and serine at positions - 141, 203, 235. We reason that the abundance of glutamic acid could result in the decreased affinity of p53 to DNA due to the local change in the ionic charge at the site of the amino acid p53 variant. Indeed, the PROVEAN tool predicted a deleterious effect on p53 function for glutamic acid at position 128. In addition, according to PHANTM classifier, C141S substitution leads to a decrease of p53 transcription activity by 41.08% as compared to wtp53.

**Figure 7:**
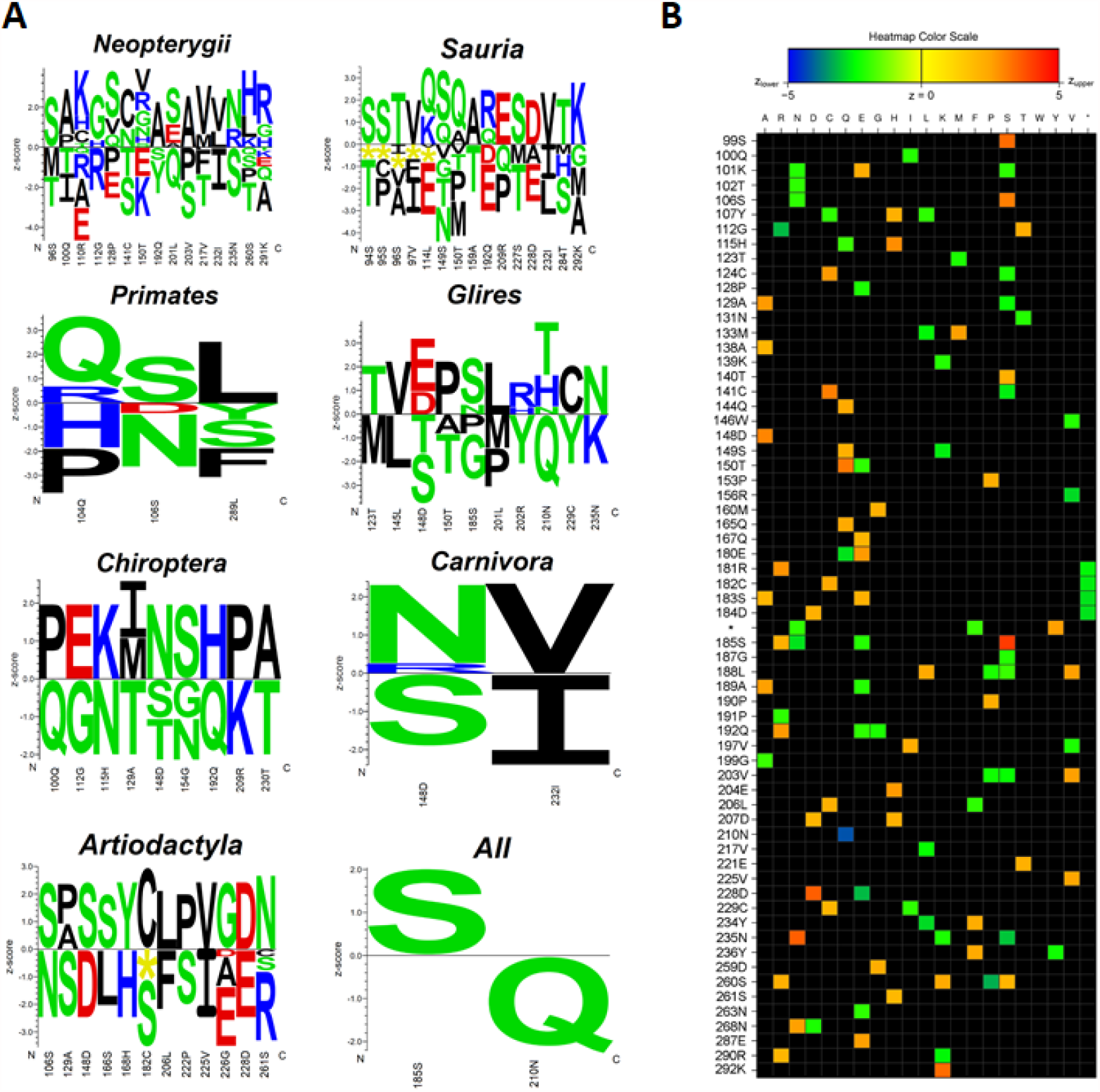
Correlation of particular p53 amino acid residues with the maximal lifespan of species. **(A)** Logos quantifying the strength of p53 core domain residue association (related to the human aa 94 – 293 according to p53 canonical sequence) with the maximal lifespan in years in the analyzed subgroups of animals. Amino acid residues on the positive y-axis are significantly associated with the prolonged lifespan phenotype and residues on the negative y-axis are significantly associated with the shorter lifespan phenotype (significance threshold p-value ≤ 0.05). The height of each letter representing the strength of the statistical association between the residue and the data set-phenotype. The amino acids are colored according to their chemical properties as follows: Acidic [DE]: red, Basic [HKR]: blue, Hydrophobic [ACFILMPVW]: black and Neutral [GNQSTY]: green. **(B)** Heatmap visualisation of the strength of residue association (without Bonferroni correction). The color-scale ranges from blue (z < -5) to red (z >5). Each column corresponds to one of the 20 proteinogenic amino acids and each row to a position in the submitted multiple sequence alignment (Supplementary Material 4).

The lifespan of species in **Sauria** is significantly variable. The organisms with the longest lifespan in this group are three-toed box turtle (*Terrapene carolina triunguis* (138 years)) and kakapo (*Strigops habroptila* (95 years)). Green anole (*Anolis carolinensis)* has the shortest lifespan in the group (7.2 years). The correlation analyses show that similarly to **Neopterygii** a specific fifteen amino acid residue fragment in the p53 core domain is associated with the prolonged lifespan (Figure 7A). The most common p53 variation for long-lived Sauria is similar to Neopterygii and it is the presence of serine (at positions 94, 95, 149 and 227 – corresponding to human p53) and the presence of valine (at positions 97 and 232, identical to Neopterygii). When compared to human p53, in short-lived organisms we identified threonine at positions 94, 149, 159 and 227 and glutamic acid at positions 114, 192 and 228. In addition, deletions in the p53 sequence were found at positions 94-97 and 114 (Figure 7A). Similar to Neopterygii, the most common p53 variation for short-lived Sauria is the presence of threonines and glutamic acid residues. However, more studies are needed to elucidate the functionality of these p53 sequences.

The organisms with the longest lifespan in **Primates** group are human (*Homo sapiens* (122 years)) and western gorilla (*Gorilla gorilla* (60 years)). Tarsier (*Carlito syrichta)* has the shortest lifespan in the group (16 years). The correlation analyses show that the specific amino acid triad, Q^104^, S^106^, L^289^ is significantly associated with a prolonged lifespan (Figure 7A). Beside a serine residue at position 106, two others; glutamine at position 104 and leucine at position 289 are both hydrophobic and might impact the structure of the DNA binding domain.

On the contrary, proline or histidine at position 104, asparagine at position 106 and phenylalanine, serine or tyrosine at position 289 are associated with short-living primates. While studying human longevity, one needs to consider that prolonged lifespan of *Homo sapiens* is associated with the cultural and socio-economical advantages. Therefore, we have additionally performed analyses upon excluding *Homo sapiens* from the dataset. The same variations were observed in the correlation analyses. Taken together, our results demonstrate that the amino acid variations shown in Figure 7A are conserved in the following closely related species: *Homo sapiens, Pan troglodytes* and *Gorilla gorilla*.

The dataset of **Glires** contains seventeen species with lifespans ranging from 3.8 to 31 years. The organisms with the longest lifespan in this group are *Heterocephalus glaber* (31 years) and *Castor canadensis* (23). The shortest lifespan in the group is *Rattus norvegicus* (3.8 years). The correlation analyses show that ten amino acid residues are significantly associated with prolonged lifespan (Figure 7A). Two threonine residue variations (positions 123 and 210) are present in long-lived Glires. Other animo acid changes occur only once. Interestingly, in short-lived Glires, there is a significant presence of threonine also at two other locations (positions 148 and 150). Similar variations are also observed in the methionine residues (at positions 123 and 201), tyrosine (positions 202 and 229) and proline (positions 185 and 201).

The organisms with the longest lifespan in the dataset of **Chiroptera** are Brandt’s bat (*Myotis brandtii* (41 years)) and little brown bat (*Myotis lucifugus* (29 years)). Pale spear-nosed bat (*Phyllostomus discolour)* has the shortest lifespan (9 years). The correlation analyses show that nine amino acid residues are associated with significant prolonged lifespan association (Figure 7A).

The organisms with the longest lifespan in **Carnivora** group are polar bear (*Ursus maritimus)* and panda (*Ailuropoda melanoleuca)*. The shortest lifespan in the group is ferret (*Mustela putorius furo*). The correlation analyses show that two amino acids in the p53 core domain, positions 148 and 232, are significantly associated with lifespan (Figure 7A). While the presence of asparagine at position 148 and valine at position 232 is associated with long lifespan, the presence of serine at position 148 and isoleucine at position 232 is associated with short lifespan.

The organism with the longest lifespan in **Artiodactyla** group is bowhead whale (*Balaena mysticetus* (211 years)) followed by orca (*Orcinus orca* (90 years)). Correlation analyses show that twelve amino acid residues in the p53 core domain are significantly associated with prolonged lifespan (Figure 7A). Similar to Neopterygii and Sauria, the most common variation present in the long-lived organisms are associated with serine at positions 106, 148 and 166 – corresponding to human p53. The variation of serine at positions 129, 182 and 222 is the most common variation for short-lived Artiodactyla, together with variation in the glutamic acid residues at positions 226 and 228.

Next, we investigated all 118 RefSeq p53 sequences to evaluate associations between amino acid variations and maximal lifespan (Figure 7A). When applying the Bonferroni correction, only two significantly associated residues were revealed, corresponding to human serine 185 and asparagine 210. Organisms that have serine at position 185 live statistically longer than organisms with another amino acid in this position. Interestingly, pS185 variants are rare in humans and only few variants have been found to be cancer-specific suggesting that S185 might be a conserved amino acid critical for organismal longevity [54]. On other hand, organisms that contain glutamine instead of asparagine at position 210 have significantly shorter maximal lifespan. Without Bonferroni correction, from the 200 analyzed positions of the aligned p53 core domains (related to human 94-293 aa), 64 positions were significantly associated with lifespan (Figure 7B). Positive correlations with longevity are shown by orange and red colors, green and blue show negative correlations. However, more detailed studies are needed to fully apprehend the functionality of the changed p53, both in the short-lived and in the long-lived organisms.

The changes at the molecular level are often a result of the adaptation of species to the environmental forces. To evaluate if the amino acid residues in the p53 core domains (aa 94 – 293 of the human p53 canonical sequence) share some relevant features in relation to the convergent evolution, we constructed a sequential circular representation of the multiple sequence alignments and the mutual information it contains (Figure 8). This figure shows that the amino acid residues significantly associated with aging (extracted from the heatmap (Fig. 7b), highlighted in light green) very often coevolved together (represented by connected lines). This observation may provide the evidence for the convergent evolution of p53 proteins in organisms with extreme longevity. According to Passow and colleagues, taxa with evidence of positive selection in the *TP*53 gene are those with the lowest incidences of cancer reported in amniotes (elephants, snakes and lizards, crocodiles and turtles) [55].

**Figure 8:**
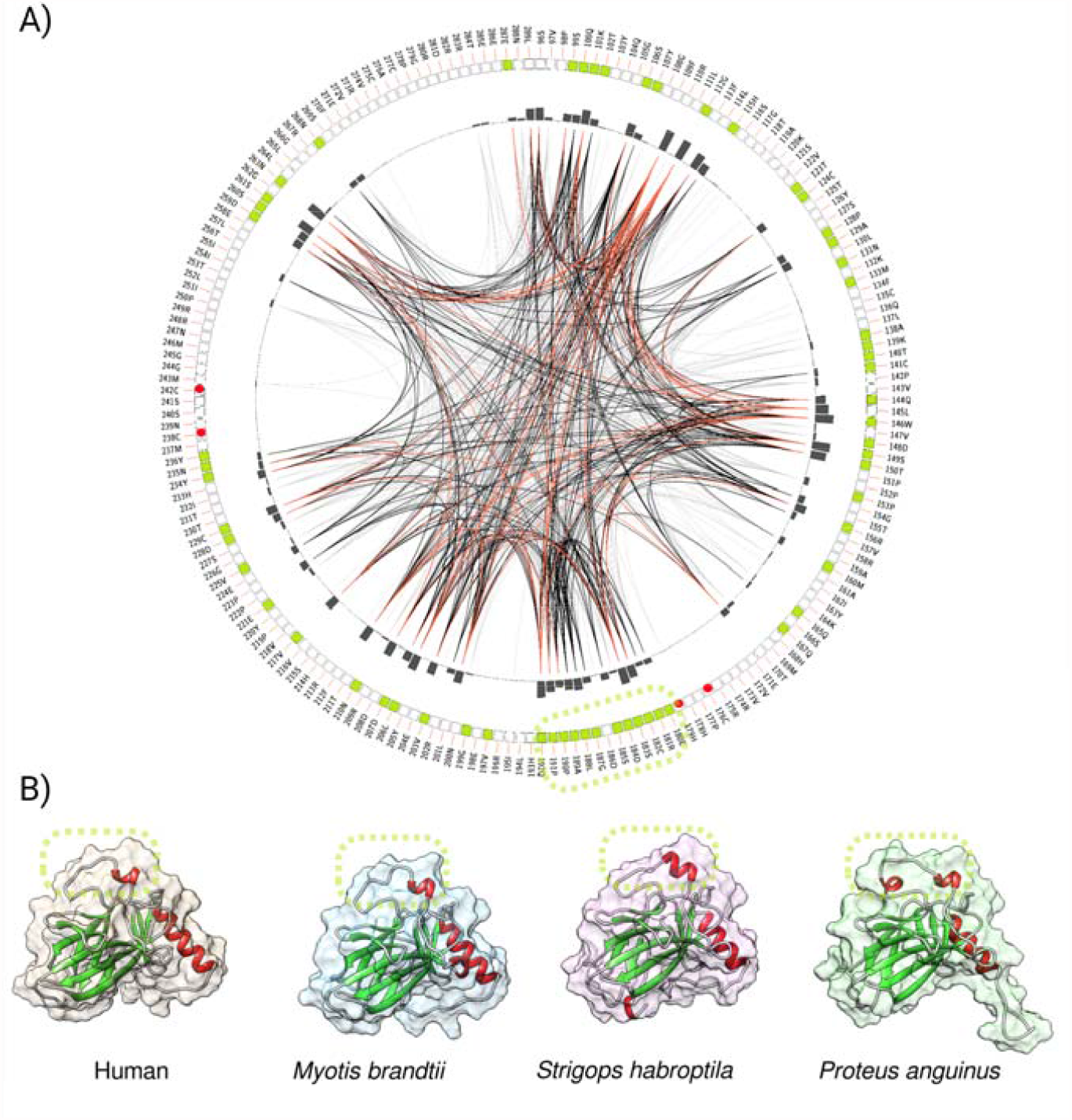
A graphic representation of the depicted amino acid changes in p53 linked to longevity in the animal kingdom. (A) Mutual information to infer convergent evolution of p53 core domains. Circos plot is a sequential circular representation of the multiple sequence alignment and the information it contains. Green boxes in the outer circle indicate amino acid residues significantly correlated with maximal lifespan, four red points indicate the Zn^2+^ binding residues. The dashed oval highlights the longest region associated with longevity. Lines connect pairs of positions with mutual information greater than 6.5 [51]. Red edges represent the top 5%, black is between 70% and 95%, and gray edges account for the remaining 70%. (B) p53 core domains of three different, long-lived organisms compared to human, modeled by trRosetta. The dashed ovals highlight the longest region associated with longevity, including S185.

Table 1 lists the species of the extreme longevity with the associated p53 variations identified in our study. Apart from the unique substitutions (*Strigops habroptila, Balaena mysticetus*) and insertions (*Myotis Brandtii, Myotis lucifugus, Proteus anguinus*), a complete lack of p53 mRNA expression was found in *Turritopsis sp*.

To gain a better inshigt into the putative changes in the p53 regulatory pathways in the long-lived species in which p53 protein remains unaltered, we analyzed the sequence of p53 regulators. Sirtuin 1 (SIRT1) deacetylates p53 in a NAD^+^-dependent manner and inhibits p53 transcription activity [56]. We found that SIRT1 has an atypical protein sequence in *Cebus imitator*, a model organism to study extreme longevity in primates, and the amino acid sequence is different from all other primate SIRT1s. 3D modeling revealed that the predicted structure of *Cebus imitator* SIRT1 is significantly different from structures of SIRT1 from *Homo sapiens* and *Sapajus apella* (a close relative of *Cebus imitator*) (Figure 9).

**Figure 9:**
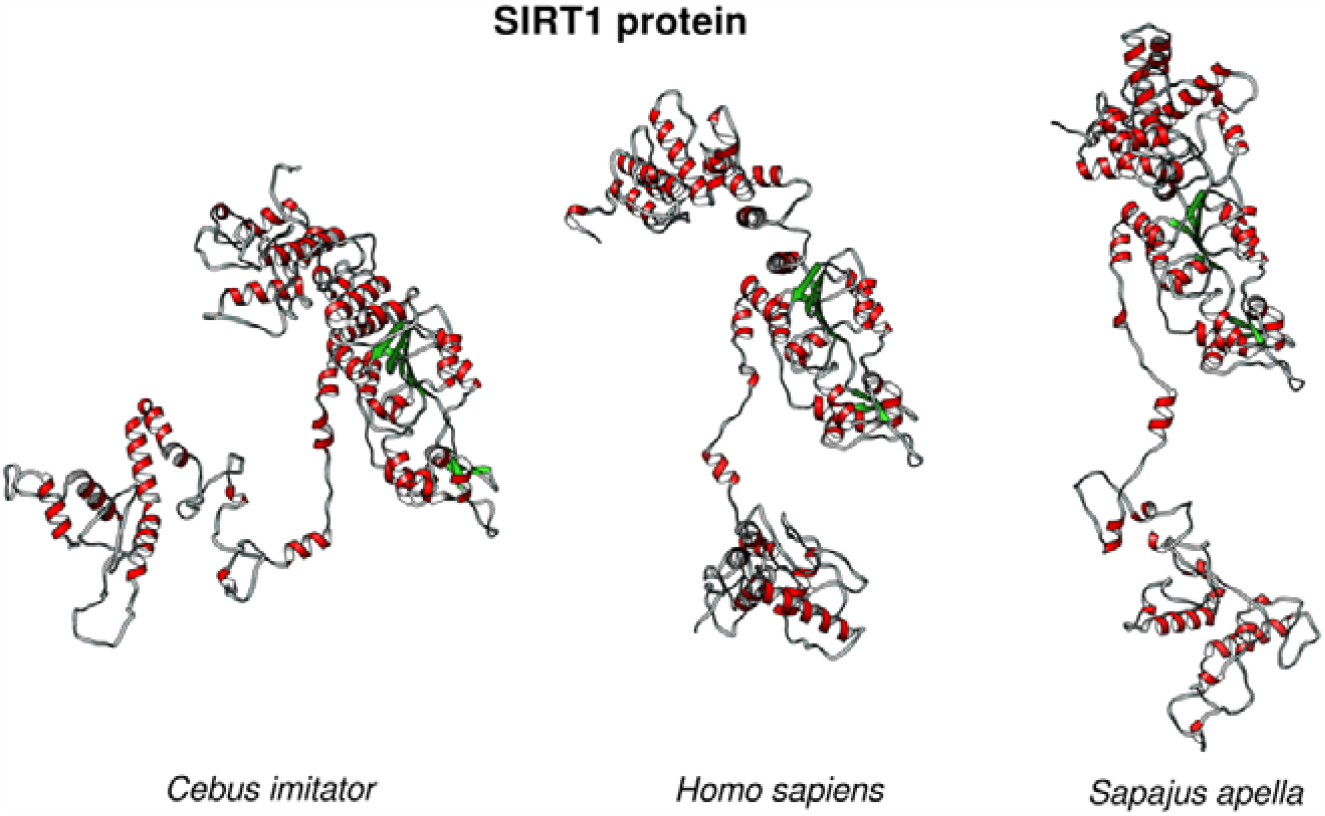
3D protein structure of SIRT1. SIRT1 structures from *Cebus imitator* (XP_017357564.1), *Homo sapiens* (NP_001135970.1) and *Sapajus apella* SIRT1 (XP_032108492.1) show differences in the protein structure in the given species.

We hypothesize, that SIRT1 from *Cebus imitator* gains new functions which might result in the decreased activity of p53, when compared to other primates, and slow down the ageing processes, most likely, by transient inhibition of p53. Yet, it remains to be elucidated which factors might be affecting the altered target-recognition by SIRT1. In addition to SIRT1, we have investigated other key factors in the p53 pathway. Suprisingly, we found that *Myotis Brandtii* (long-living bat described above), in addition to p53, has two atypical protein sequences, one in UFM1 (Ubiquitin-fold modifier 1, XP_005862786.1) and other in the p73 (tumor protein 73, XP_014401672).

Recently, it has been reported that UFM1 covalently modifies p53 and this phenomenon is called UFMylation [57]. UFMylated p53 is stabilized on the protein level as this colavalent modification antagonizes p53 ubiquitination and proteasomal degradation.

In UFM1, we found approximately 20-amino-acid-long extension of the C-terminal end in *Myotis brandtii* (and also in two other Myotis bats – *Myotis lucifugus* and *Myotis myotis*), see Figure 10. On the contrary, in other bats that live much shorter (e.g. the closest Myotis bats relative is *Pipistrellus kuhlii* with the maximal lifespan of only 8 years) and in the rest of mammals including humans, no such an extension occurs. We hypothesize that the extended UFM1 protein might contribute to the extreme longevity in Myotis bats through the loss of function and consequent p53 protein degradation. Yet, more experimental evidence is needed to draw a clear conclusion.

**Fig. 10.**
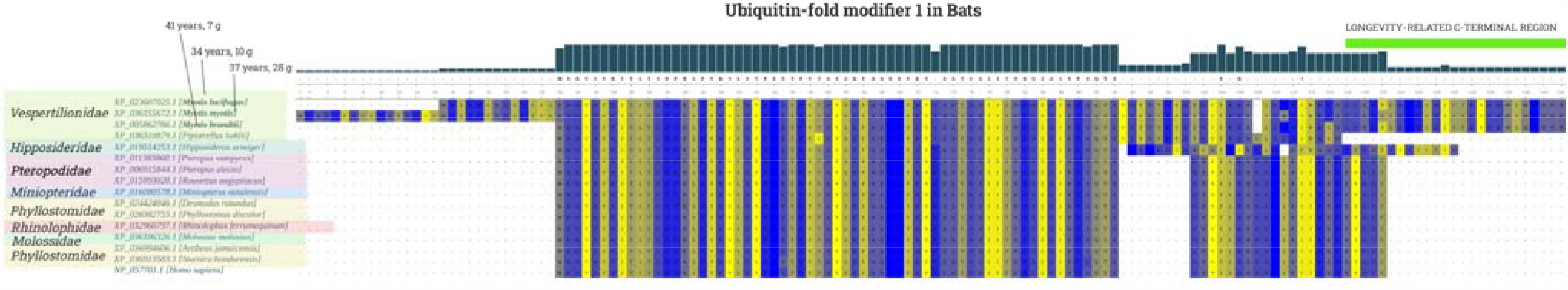
Sequence aligment of UFM1 proteins in the sequenced bat species and human. 20-amino-acid-long extension of the C-terminal end in three long-living bats is depicted in light green. Multiple sequence protein alignments of UFM1 reference protein sequences was performed in MUSCLE with default parameters [47], colors express strand propensity.

Lastly, we found the unique changes in p73 protein sequence in the *Myotis Brandtii* bat (XP_014401672). There are multiple large deletions (>10 amino acid residues) of critical p73 regions, found exclusively in this extreme long-lived bat. p73 is the transcription factor which undergoes similar cellular regulation as p53 protein and its role in aging is attibuted to induction of senescence though upregulation of CDKN1 [58].

Taken together, our analyses revealed unexpected correlation between p53 sequence variations and longevity in the animal kingdom. The changes may affect p53 functionality and thus, influence the activation of replicative senescence. In long-lived species, with no changes of p53, the upstream regulatory proteins display amino acid changes that my affect their funcitonality and that includes SIRT1 and UFM1. Yet, further studies are needed to fully comprehend the role of amino acid changes in p53 and its role in long-lived species described in our work.

## 3. Discussion

The p53 protein is a well-known tumor suppressor and the *TP*53 gene is the most often mutated gene in human cancers. On the cellular level, in humans, decreased p53 functionality is essential for cells’ immortalization and neoplastic transformation [59]. However, the role of variations in the p53 amino acid sequence at the organism level in other animals has not been studied systematically. Here, we presented an in-depth correlation analysis manifesting the dependencies between p53 variations and organismal lifespan to address the role of p53 in longevity in the animal kingdom. To date, p53 expression has been detected in all sequenced animals from unicellular Holozoans to vertebrates [32]. The seminal work by Kubota provided important evidence demonstrating that immortality is not just a hypothetical phenomenon. He demonstrated that Cnidarian species *Turritopsis* jellyfish is immortal and can repeatedly rejuvenate, reverse its life-cycle and, thus, was the first and only known “immortal” animal on earth [60]. Here, we inspected recently published data from the whole-transcriptome data of “immortal” *Turritopsis sp*. [61] and surprisingly found no expression of any of the p53 family members in the pooled data from all individuals at all developmental stages (polyp, dumpling with a short stolon, dumpling and medusa). This points to the possibility that the absence of p53 in *Turritopsis* might be directly related to its unique ability of the life cycle reversal and “immortality”.

Telomere shortening in humans induce replicative senescnence, a process regulated by p53. In the absence of p53 the replicative lifespan of human cells is extended and the concurent loss of pRB extends the replicative lifespan to a greater extend (reviewed in [62].

Intriquingly, our results obtained by using Protein Variation Effect Analyzer [63] show that the variability in lifespan among closely related species correlates with specific p53 variations. Long-lived organisms are characterized by in-frame deletions/changes, insertions or specific substitutions in the p53 amino acid sequence. It is likely that the changes imposed on p53 in long-lived species enable p53 to interact with different multiple protein partners to induce gene expression programmes varying from those induced in species with relatively normal lifespan. Based on what is know about the processes underlyign aging, we can anticipate that these gene expression programmes would enable the following changes (Figure 11): 1. more efficient tissue repair through autophagy, 2. loss of replicative senescence, 3. enhanced clearance of senescent cells by the immune system, 4. enhanced regulation of intracellular ROS levels 5. improved resistance of mitochondria to ROS-induced damage or 6. loss of immune senescence that occurs in humans during healthy aging. All of the above processes have been previously described as significantly contributing to longevity in humans (reviewed in [18]). Intriguingly, a recent GWAS study on 1 million parent lifespans, identified only several variants influencing lifespan at genome-wide significance including *CDKN2B-AS1* and *IGF2R*. The *TP*53 gene was not among the singled-out variants which, in accordance with our observations, indicate that no changes in human p53 might be attributing to longevity in humans [64]. Our analysis demonstrates that the long-lived organisms might have different, mechanisms of protection against cancer not directly linked to p53 activity. We speculate that their lifespan is not limited by somatic cells’ senescence caused by chronic stress-induced hyperactive p53 protein, which is the case for other species with shorter lifespan (Figure 11).

**Figure 11.**
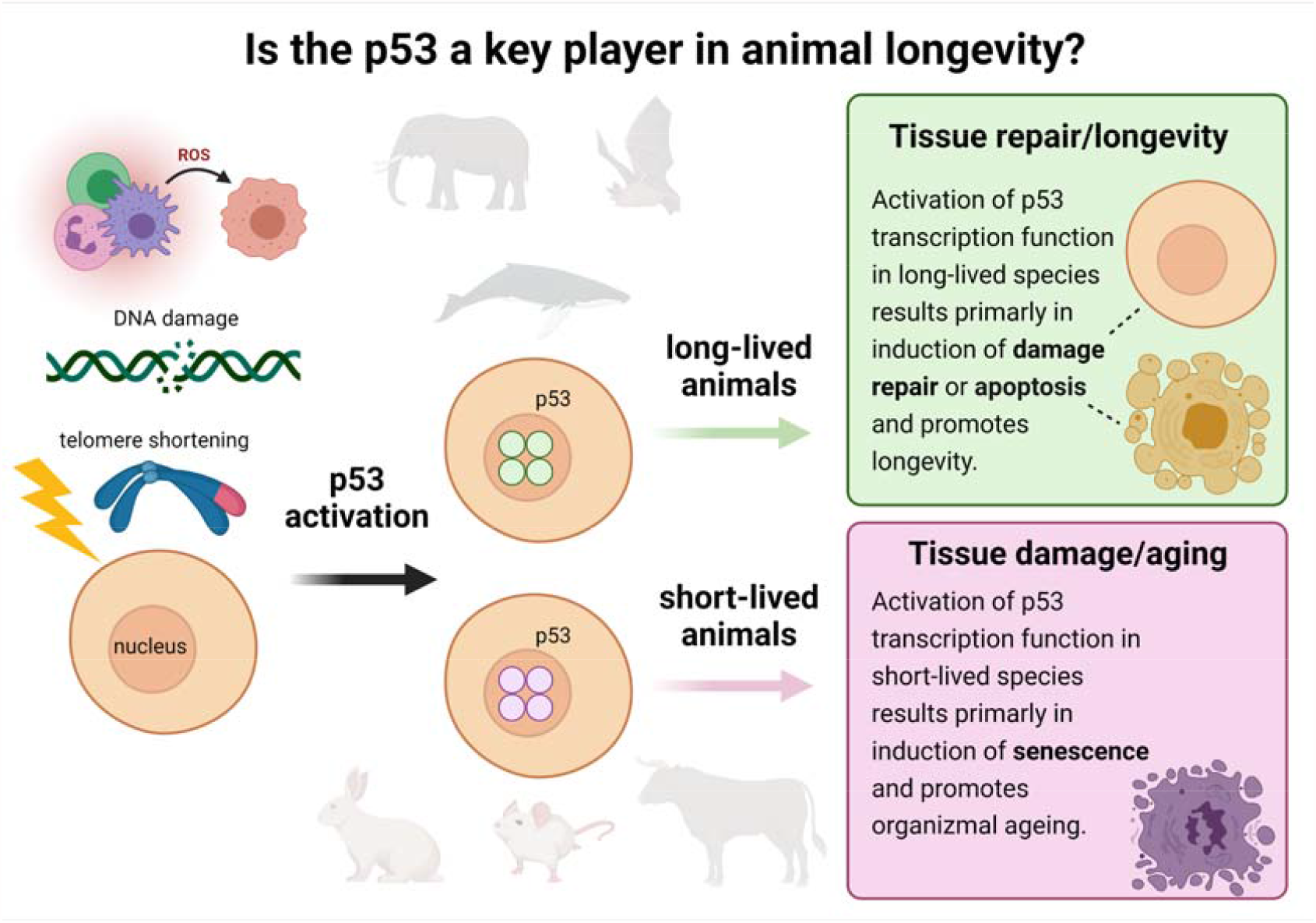
Proposed p53-centric theory of extreme longevity. Cell damage caused by ROS, DNA damage, telomere shortening, or other factors activate p53 to enable DNA repair and/or activation of the apoptosis pathway. On the other side, high activity of p53 promotes organizmal ageing, thus shortening the lifespan. We hypothesize that long-lived animals developed the ‘improved’ p53 proteins, that are less active than in their short-living counterparts, but still may sufficiently contribute to DNA damage repair in species exposed to environmental genotoxic stresses.

The maximal lifespan according to the AnAge database is attributed to Greenland shark with an estimated maximal life span of 300–500 years. Unfortunately, no transcriptomic nor genomic data for the Greenland shark (*Somniosus microcephalus*) are available. Compared to other sharks (with life expectancy of up to seventy years), its lifespan is exceptional. It will be thus very interesting to know the sequence of their p53 protein. A recent study suggest that certain animals may have evolved to have longer lifespans compared to other species belonging to the same taxa [65]. In addition, the authors found that the outliers among taxa (in terms of maximum age) have always longer lifespans, not shorter. This would support our hypothesis, that extreme longevity is a result of adaptive mutational changes in the particular critical gene(s) allowing the organisms to escape the senescence machinery.

Several experimental data support our hypothesis that specific p53 variation associates with longevity. For example, it has been found that the reduced expression of the *Caenorhabditis elegans* p53 ortholog, cep-1, results in increased longevity [66]. It has also been demonstrated that neuronal expression of p53 dominant-negative proteins, inhibiting the function of full length p53, in adult *Drosophila melanogaster* extends lifespan [67]. The same principle is most probably present in humans, where, for example, p53 variants predisposing to cancer are present in healthy centenarians [68] and a meta-analyses showed that the codon 72 polymorphic variant of p53 with proline (compared to arginine) provides an increased cancer risk, but increased survival [69]. In a recent study by Zhao et al., polymorphism at position 72 (P72 compared to R72) was reported to have a positive effect on lifespan and to delay the development of aging-related phenotypes in mice, supporting a role of the changed p53 activity in longevity [70]. Another example of a long-lived vertebrate is the elephant, which has 20 copies of the *TP53* gene [71]. In this specie, part of the DNA-binding region of p53 is deleted in all but one of the *TP*53 gene copies, which may result in the formation of dysfunctional p53 tetramers, thus, presumably, modulating p53 transcriptional activity in response to stress [71]. On the contrary, a study by Tejada-Martinez et al. [72] in cetaceans didn’t identify *TP*53 as a gene associated with extreme longevity. Nonetheless, the authors provided the evidence that natural selection in tumour suppressor genes (i.e. also in *TP*53) could act on species with extended lifespan.

It is also worth mentioning, that the animals with an extreme longevity that we and others identified, are mostly nocturnal or live in absolute darkness. These include *Strigops habroptila, Myotis brandtii, Myotis lucifugus, Proteus anguinus, Balaena mysticetus, Heterocephalus glaber*. It is thus, likely, that low or no exposure to UV-induce damage promotes the evolutionary changes in p53 protein structure that alters p53 activity. Also, in such species, the endogenous levels of reactive oxygen species might be lower when compared to animals from other, less extreme habitats. This might be a consequence of the changes in the metabolism rate that might affect the overall rate of oxidative phosphorylation and in consequence slow down generation of free radicals through electron transport chain. This hypothesis is in agreement with the recent study showing that rapamycin, a widely studied inhibitor of mTOR, prevents UV-induced skin ageing through inhibition of p53, reversal of UVA-induced cellular senescence and induction of authophagy [73].

Despite high complexity of the p53 proteins family, modern methods of comparative genomics provide useful tools to explore protein variations in closely related species, and to correlate the extracted molecular information with lifespan [74]. According to Sahin and DePinho, the hyperactivity of p53 in the presence of accumulated DNA damage and ROS is one of the main causes of aging [41]. This observation is in congruence with our hypothesis that organisms with atypical p53 sequences, likely attenuating the wtp53 activity, are extremely long-lived. Of note, even if several p53 amino acid changes have been found in various animal groups, some variations developed in convergent evolutions in different groups of species. For example, the presence of threonine and glutamic acid was observed in short-lived organisms of different groups and richness of serine residues was typical for long-lived organisms in several groups and serine residue at position 185 is significantly associated with prolonged lifespan across all analyzed species. Yet, further mechanistic studies are needed to pin down how certain changes in p53 affect its functionality and how they contribute to longevity.

## 4. Materials and Methods

### 4.1. Searches of maximal lifespan

To access data of longevity and maximal lifespan, we used AnAge Database (https://genomics.senescence.info/species/), AnAge currently contains data on longevity of more than four thousand animals.) [45]. We have downloaded the whole dataset and selected species presented in NCBI RefSeq database.

### 4.2. Protein similarity searches

For protein similarity searches we have downloaded all available p53 sequences from RefSeq database (https://www.ncbi.nlm.nih.gov/refseq/) and merged them with AnAge Database. We received the p53 sequence information of 118 species with information about their lifespan and sorted them according to their phylogenetic group (Supplementary Material 1). In animals with extreme longevity, where the p53 homologs were not present in NCBI, local blast searches (tblastn) applied on *de novo* assembled transcriptomes were used together with the default “BLAST+ make database” command and searching parameters within UGENE standalone program [75].

### 4.3. Transcriptome assemblies

Transcriptomic data for bowhead whale was obtained from http://www.bowhead-whale.org/ [46]. When there were only raw seq reads from the RNA-seq experiments available (deposited in the NCBI SRA), we performed the *de novo* assembly first, using Trinity tool [76] from the Galaxy webserver (https://usegalaxy.eu/)[77] with default settings. This was done for *Proteus anguinus* (SRX2382497) and *Sphenodon punctatus* (SRX4014663), resulting assemblies are enclosed in Supplementary Material 5.

### 4.4. p53 protein tree and real phylogenetic tree construction

The protein tree was built using Phylogeny.fr platform (http://www.phylogeny.fr/alacarte.cgi) [78,79] and comprised the following steps. First, the sequences were aligned with MUSCLE (v3.8.31) [47] configured for the highest accuracy (MUSCLE with default settings). After alignment, ambiguous regions (i.e. containing gaps and/or poorly aligned) were removed with Gblocks (v0.91b) [80] using the following parameters: -minimum length of a block after gap cleaning: 10; -no gap positions were allowed in the final alignment; -all segments with contiguous non-conserved positions longer than 8 were rejected; -minimum number of sequences for a flank position: 85%. The phylogenetic tree was constructed using the maximum likelihood method implemented in the PhyML program (v3.1/3.0 aLRT) [81,82]. The JTT substitution model was selected assuming an estimated proportion of invariant sites (of 0.204) and 4 gamma-distributed rate categories to account for rate heterogeneity across sites. The gamma shape parameter was estimated directly from the data (gamma=0.657). Reliability for the internal branch was assessed using the bootstrapping method (100 bootstrap replicates). Graphical representation and edition of the phylogenetic tree were performed with TreeDyn (v198.3) [83]. Real phylogenetic tree was reconstructed using PhyloT (https://phylot.biobyte.de/) and visualized in iTOL (https://itol.embl.de/) [84].

### 4.5. Prediction and statistical evaluation by PROVEAN

The effect of the p53 variations in long-lived organisms was predicted and statistically evaluated by Protein Variation Effect Analyzer web-based tool (PROVEAN; http://provean.jcvi.org/index.php) [63,85]. PROVEAN is a software tool which predicts whether an amino acid substitution or in/del has an impact on the biological function of a protein [63]. All inspected p53 variations in selected animals were statistically evaluated and numbered according to the human canonical p53 sequence (NP_000537.3).

### 4.6. Modelling of 3D Protein Structures

We used SWISS-MODEL template-based approach (https://www.swissmodel.expasy.org/interactive) [86] to predict 3D structures using individual FASTA sequences and reference PDB:4mzr as the crystal structure of p53 tetramer from *Homo sapiens* with bound DNA [87]. Resulting PDB files are enclosed in Supplementary Material 6. Predicted p53 structures were visualized in UCSF Chimera 1.12 [88]. Effects of the novel mutation on SIRT tertiary structure were predicted by RaptorX [89].

### 4.8. Correlation of maximal lifespan and alterations within the p53 core domain in vertebrates

Residue level genotype/phenotype correlations in p53 multiple sequence alignment were performed using SigniSite 2.1 (http://www.cbs.dtu.dk/services/SigniSite/) [53] with significance threshold p-value ≤ 0.05. Bonferroni single-step correction for multiple testing was applied for the global correlation of all sequences, no correction was applied for smaller groups of taxonomically related animals. The manually curated set of 118 high-quality p53 protein sequences obtained from the NCBI (https://www.ncbi.nlm.nih.gov/) was used as input file. These sequences were taken from the RefSeq database and the canonical isoform corresponding to human full length p53 isoform a (NP_001119584.1) was manually filtered for each vertebrate species. The resulting set of these 118 p53 sequences was aligned within UGENE workflow [75], MUSCLE algorithm [47] with default parameters. All sequences were then manually trimmed to preserve only the core domain, which corresponds to human 94 – 293 aa. Then the numerical values of maximal lifespan of each organism were added into the resulting FASTA file, based on the information in the reference AnAge database (http://genomics.senescence.info/species/) [45]

### 4.9. Convergent evolution

Multiple sequence alignment of p53 core domains from 118 species was uploaded to the MISTIC webserver (http://mistic.leloir.org.ar/index.php), with PDB 2ocj (A) as the reference and using default parameters [90].

### 4.10. Gene gain and losses

*TP*53 gene gain or losses were inspected using Ensembl Comparative Genomics toolshed [91] via Ensembl web pages and *TP53* gene query ENSG00000141510: https://www.ensembl.org/Homo_sapiens/Gene/SpeciesTree?db=core;g=ENSG00000141 510;r=17:7661779-7687550

## 5. Conclusions

This study reveals a previously overlooked correlation between longevity and a potential change in p53 function due to the amino acid variations across the animal kingdom. Strikingly, several long-lived species, including *Myotis brandtii, Myotis lucifugus, Balaena mysticetus, Heterocephalus glaber, Strigops habroptila* and *Proteus anguinus* display unique p53 protein sequence properties not shared with their close relatives that have a shorter lifespan. Altogether, our evidence suggests convergent evolution of p53 sequences supporting a higher insensitivity to p53-mediated senescence under prolonged stress conditions in long-lived vertebrates. Our observations that specific variations of p53 protein are correlated with lifespan provide important grounds for further exploration of p53 sequences in species displaying extreme longevity. Most importantly, our data implies a general mechanism at work in all vertebrates leading to extended lifespan, which might be translated to studies on extension of the health span in humans.

## Supplementary Materials

Supplementary material 1: Table of all analyzed organisms, their maximal lifespan and reference p53 sequences; Supplementary material 2: 3D structures of p53 core domains; Supplementary material 3: The list of organisms with the longest and shortest lifespan in particular phylogenetic groups; Supplementary material 4: Multiple sequence alignment of analyzed p53 sequences; Supplementary material 5: Transcriptome assemblies of Proteus anguinus (SRX2382497) and Sphenodon punctatus (SRX4014663); Supplementary material 6: Predicted structures of p53 homologs in PDB format

## Author Contributions

Conceptualization, M.B., V.B. and P.P.; methodology, M.B.; validation, V.B., J.C.; formal analysis, M.B., V.B., A.V.; investigation, M.B. and J.ZP.; resources, M.B., A.V. and V.B..; data curation, M.B., A.V. and J.C.; writing—original draft preparation, M.B., J.ZP., and V.B.; writing—review and editing, M.B., J.C., P.P., J.ZP.; visualization, M.B. and A.V.; supervision, P.P and J.ZP.; project administration, J.ZP.; funding acquisition, P.P. All authors have read and agreed to the published version of the manuscript.

## Institutional Review Board Statement

Not applicable. Informed Consent Statement: Not applicable.

## Data Availability Statement

The data presented in this study are available in supplementary material.

## Acknowledgments

We thank Dr. Jean-Christophe Bourdon for valuable comments and discussion, Dr. Philip Coates for proofreading and editing. We would also like to express our gratitude to M. Sc. Alena Volná and Milan Bolek for their time spent on illustration preparation. This work was supported by The Czech Science Foundation (18-15548S) “ and by the Warsaw University’s Integrated Development Programme (ZIP), co-financed by the European Social Fund, European Union Operational Programme Knowledge, Education, Development for 2014–2020 (POWER), under programme priority axis III: Higher Education for the Economy and Development, action 3.5. It is implemented on the basis of an agreement between the University of Warsaw and The National Center for Research and Development, an implementing agency of the Ministry of Science and Higher Education.

