## Supplementary material 2 for "The changes in the p53 protein across the animal kingdom pointing to its involvement in longevity"

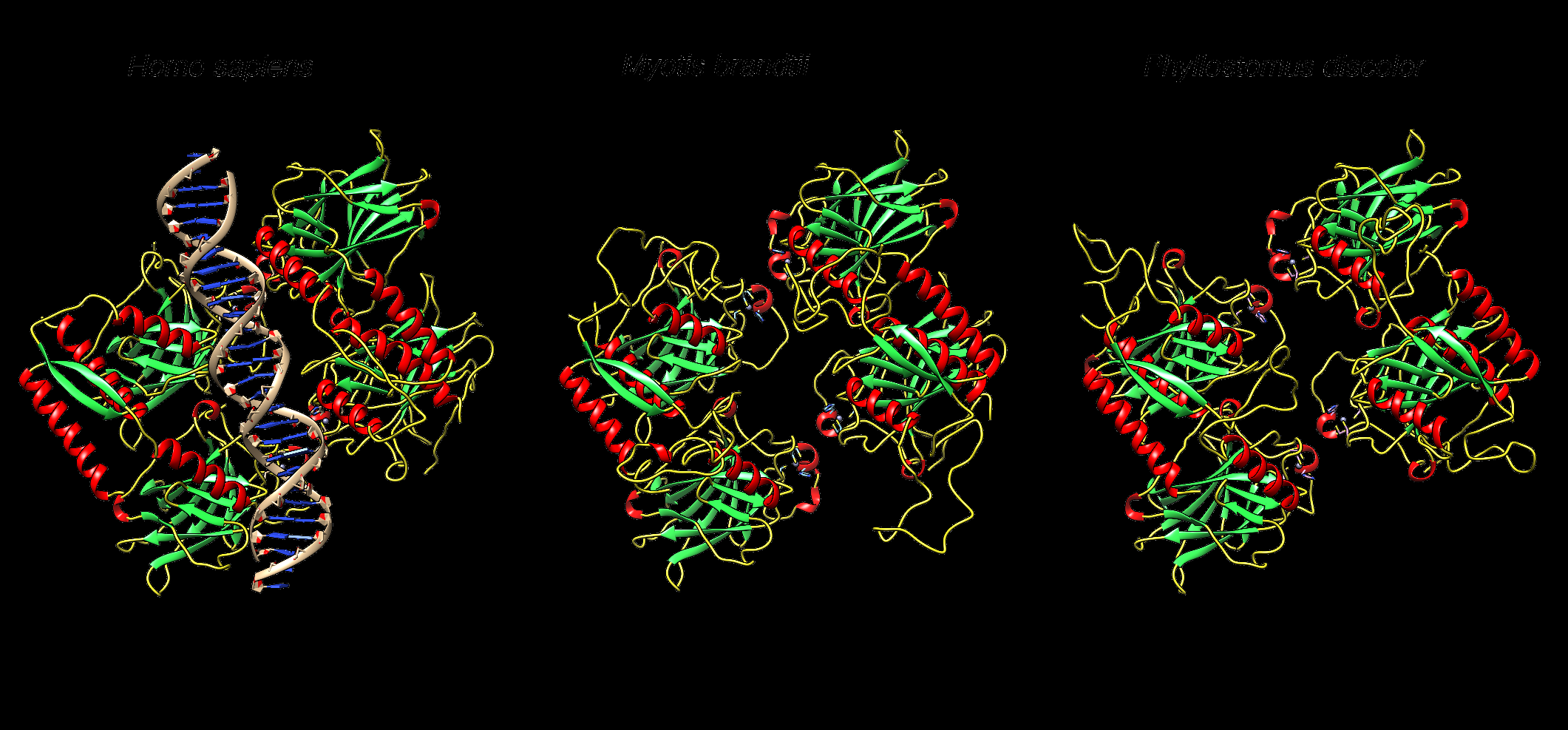

**Supplementary material 2:** Experimental (human) and modelled (bats) 3D structures of tetramerized p53 core domains. (Left) *Homo sapiens* p53 tetramer binding DNA (Middle) p53 tetramer from long-lived bat *Myotis Brandtii* (Right) p53 tetramer from short-lived bat *Phyllostomus discolor.* The insertion in the DNA binding domain of bats with long lifespan is present inside the DNA interaction cavity, suggesting decreased affinity of p53 for binding to DNA (blue circle).
