## Supplementary material 3 for "The changes in the p53 protein across the animal kingdom pointing to its involvement in longevity"

**Supplementary material 3:** The list of organisms with the longest and shortest lifespan in particular phylogenetic groups. The total number of analyzed animals for each group is depicted in column “Count”. Formal and common names and their maximal lifespans in years are given for each organism.

| **Group** | **Count** | **Shortest living animal (years)** | **Longest living animal (years)** |
| --- | --- | --- | --- |
| Afrotheria | 3 | *Echinops telfairi* (19) Lesser hedgehog tenrec | *Trichechus manatus latirostris* (69) Florida manatee |
| Artiodactyla | 13 | *Capra hircus* (20.8) Goat | *Balaena mysticetus* (211) Bowhead whale |
| Batrachia | 3 | *Ambystoma mexicanum* (17) Axolotl | *Proteus anguinus* (102) Olm |
| Carnivora | 10 | *Mustela putorius furo* (11.1) Ferret | *Ursus maritimus* (43.8) Polar bear |
| Eulipotyphla | 3 | *Condylura cristata* (2.5) Star-nosed mole | *Erinaceus europaeus* (11.7) European hedgehog |
| Glires | 17 | *Rattus norvegicus* (3.8) Brown rat | *Heterocephalus glaber* (31) Naked-mole rat |
| Chiroptera | 8 | *Phyllostomus discolor* (9) Pale spear-nosed bat | *Myotis Brandtii* (41) Brandt´s bat |
| Metatheria | 3 | *Sarcophilus harrisii* (13) Tasmanian devil | *Vombatus ursinus* (30) Common vombat |
| Neopterygii | 24 | *Betta splendens* (2) Siamese fighting fish | *Cyprinus carpio* (47) Common carp |
| Perissodactyla | 3 | *Ceratotherium simum simum* (45) Southern white rhinoceros | *Equus caballus* (57) Horse |
| Primates | 15 | *Carlito syrichta* (16) Philippine tarsier | *Homo sapiens* (122.5)  Human |
| Sauria | 13 | *Anolis carolinensis* (7.2) Green anole lizard | *Terrapene carolina triunguis* (138) Three-toed box turtle |
